# Click-xylosides overcome neurotoxic effects of reactive astrocytes and promote neuronal growth in a cell culture model of brain injury

**DOI:** 10.1101/286450

**Authors:** Swarup Vimal, Balagurunathan Kuberan

## Abstract

Astrocytes, upon activation in response to brain injury, play a critical role in protecting neurons by limiting inflammation through the excessive secretion of many soluble factors, such as, chondroitin sulfate proteoglycans (CSPGs). Unfortunately, excessive CSPGs paradoxically prohibit neuronal recovery and growth, and eventually constitute a scar tissue. Many studies have attempted to overcome this barrier through various molecular approaches including the removal of inhibitory CSPGs by applying chondroitinase enzymes. In this study, we examined whether click-xylosides, which serve as primers of glycosaminoglycan (GAG) biosynthesis, can compete with endogenous inhibitory CSPGs for GAG assembly by serving as decoy molecules and thereby potentially reverse reactive astrocyte mediated neuronal growth inhibition. We investigated the axonal growth of hippocampal neurons in the presence of xyloside treated and untreated reactive astrocyte-conditioned media as a model recapitulating brain injury. Click-xylosides were found to interfere with the GAG biosynthetic machinery in astrocytes and reduced the amount of secreted inhibitory CSPGs by competing with endogenous assembly sites. The extent of underglycosylation was directly related to the outgrowth of hippocampal neurons. Overall, this study suggests that click-xylosides are promising therapeutic agents to treat CNS injuries and warrants further in vivo investigations.

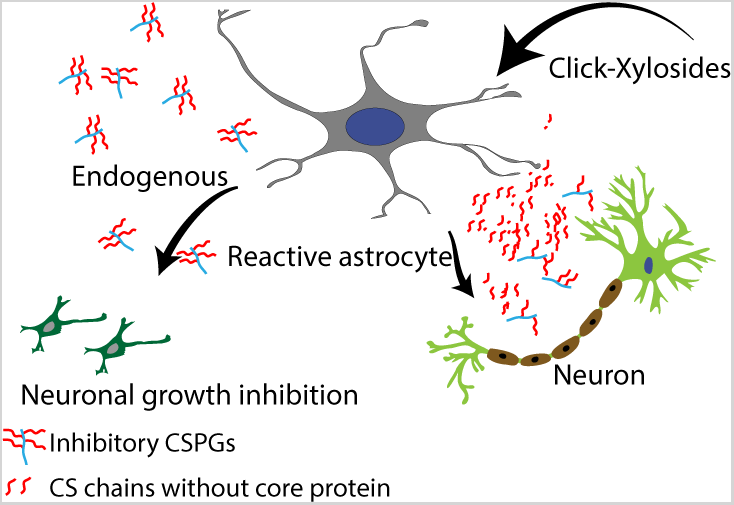

---

Central Nervous System (CNS) lesions cause reactive behavior in glia, leading to the formation of a glial scar that is beneficial immediately following an injury but then turns to become detrimental to axonal growth/regeneration in later stages. By presenting inhibitory cues, such as chondroitin sulfate proteoglycans (CSPGs), the scar tissue suppresses regeneration and growth of the damaged neurons ^1^.

CSPGs are composed of multiple CS glycosaminoglycan (GAG) side chains attached to a core protein and are known to play a dominant role in neuronal inhibition. Although their mechanistic pathway is only partially understood, CSPGs have been shown to interact with various cell surface receptors found on axons, resulting in cytoskeletal changes followed by the retraction of neurites from the scar site ^2^. The neurotoxic effects of the reactive astrocytes have been attributed to the CS side chains of the CSPGs. Therefore, various regenerative therapies have focused on the removal of the CS side chains from inhibitory CSPGs to facilitate neuronal growth ^3,4^. Chondroitinase ABC (ChABC)-mediated digestion and removal of CS side chains has been, so far, the most promising approach to minimize neuronal growth inhibition from the glial scar ^3^. Cua and Lau recently described another approach by using metalloproteases that cleave the core protein of CSPGs instead. They used ADAMTSs to reduce or abolish the inhibition of neurite outgrowth mediated by purified inhibitory CSPG or reactive astrocyte-derived ECM^5^. However, there are significant limitations in the use of enzyme-based therapeutic approaches to remove CSPGs in the clinical setting. First, inhibitory CSPGs are secreted in a sustained manner over a long period after an injury and therefore, a single interventional dosage of ChABC, which has a short half life *in vivo*, ^6^ may not be sufficient. In addition, some CS variants exhibit growth-promoting roles in the CNS. Therefore, indiscriminately removing all CSPGs might be counterproductive and hinders CNS recovery in the long term. Other approaches, such as the use of fluorinated glucosamine or SiRNA, have been studied to hinder GAG biosynthesis^11-13^

An alternative approach to CSPG degradation could be the use of small molecules to reduce the degree of glycosylation (attachment of CS side chains) of the core proteins. CSPG biosynthesis is initiated by the covalent attachment of a xylose monosaccharide to select serine residues on the core protein. Subsequently a tetrasaccharide linkage region is assembled and then extended by sequential addition of hexosamine and glucuronic acid residues to produce CS chains ^7^. Synthetic xylosides are small molecules in which xylose molecules are attached to various hydrophobic aglycones at the anomeric carbon. Exogenously added xylosides can compete with endogenous xylosylated core proteins in astrocytes and hence affect CS attachments to the core protein in terms of chain length, sulfation pattern, and the number of CS chains. In fact, β-D-xylosides have been shown to down regulate CSPGs production in embryonic chick cells ^10^. Other approaches, such as the use of fluorinated glucosamine or SiRNA, have been studied to hinder GAG biosynthesis^11-13^.

The use of β-D-xylosides to alter proteoglycan synthesis in astrocytes *in vitro* has been demonstrated. However, the glycosidic bond of previously used xylosides is inherently unstable and β-D-xylosides are only effective at very high concentrations of 1 to 5 mM, which are unsuitable for therapeutic uses ^14^. Our lab previously employed click-chemistry to attach various aglycone residues to xylose through a triazole ring to synthesize click-xylosides. These molecules have increased metabolic stability and hence enhanced intra-cellular availability in the endoplasmic reticulum and the golgi apparatus, the sites of CS biosynthesis ^12^. Since aglycone residues affect the quantity of GAGs produced ^13^, we hypothesized that by varying the aglycone, one can modulate the CS attachments present in the core proteins of CSPGs, through competing with precursors for GAG assembly, hence limiting the inhibitory potential of CSPGs that typically contain between two to four CS chains per protein ^14^.

To evaluate the effect of click-xylosides on inhibitory potential of astrocyte reactive media, primary rat cortical astrocytes were treated with 100 µM click-xylosides for 48 hours. Next, click-xyloside-treated conditioned media was collected and added to a 24-hour-old culture of embryonic day 18 primary rat hippocampal neurons (see materials and methods for details). The structures of click-xylosides used in the study are shown in Fig 1. The representative growth of neurons from click-xyloside treatments (and controls) is shown in Fig 2. The neuronal outgrowth was minimal in CM harvested from the reactive astrocyte untreated with xylosides and maximal in CM harvested from XY3-and XY4-pre-treated reactive astrocytes. In fact, the growth response to XY4-treated CM was similar to neurons treated with fresh F12 media (Fig 2 (d) and (f)).

**FIGURE 1:**
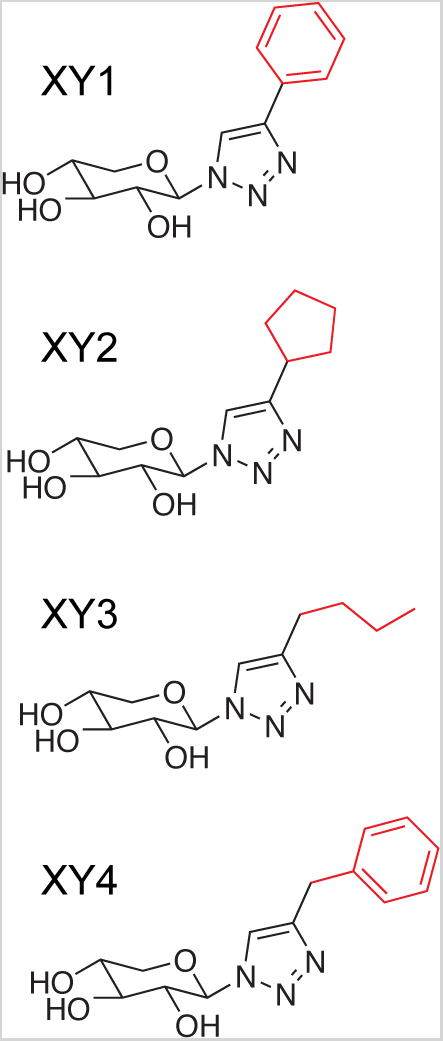
Structures of click-xylosides used in the study.

**FIGURE 2:**
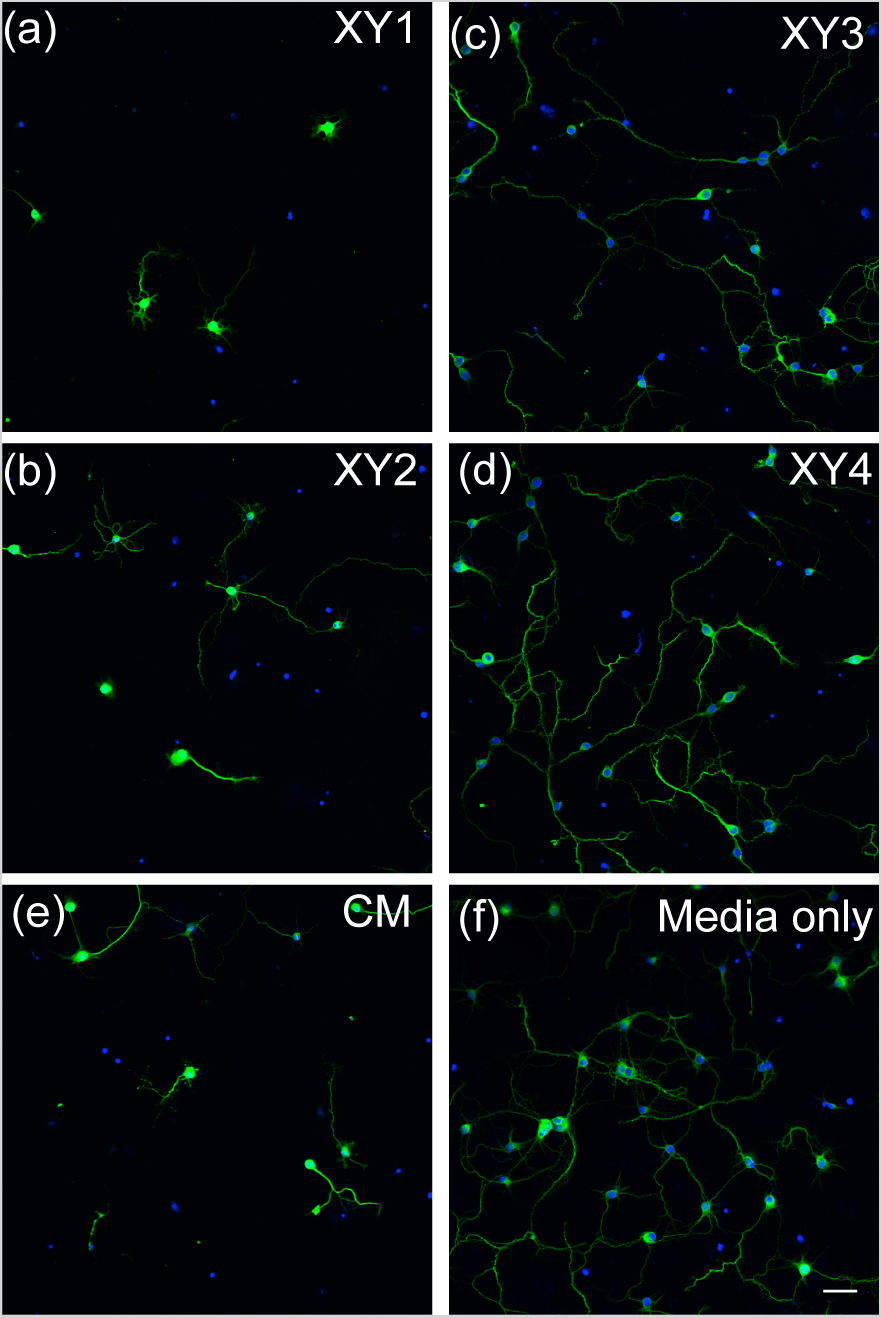
Neuronal growth responses to various click-xyloside treatments. (a-d) Immunostained images of neurons treated with CM from XY_1_, XY_2_, XY_3_, and XY_4_-pre-treated astrocytes, respectively. (e) and (f) show outgrowth of neurons treated with control CM and fresh media respectively. Scale bar = 50 μm, green: anti-rat TAU/MAPT, blue: DAPI.

We used Calcein AM dye staining to quantify the neuronal response to click-xyloside treatments. As shown in Fig S1, fluorescence signals from Calcein AM-treated neurons were found both in cell bodies and neurite outgrowths. Therefore, the fluorescence read outs quantify both neuronal viability and outgrowth. Fig 3(a) compares the relative influence of various click-xyloside-primed CM on neurons, normalized with signal intensity from CM treated neurons. The bars quantify the outgrowth behavior shown in Fig (2). Viability and outgrowth in CM from XY4 treatment were comparable with those of untreated neurons, suggesting that XY4 reverts the inhibitory potential of astrocytes (p value of fresh media vs. XY4 treatments was 0.474). In contrast, no change in neuronal viability or outgrowth was noted for neurons treated with XY1-primed CM compared with control CM treatments (p value of CM vs. XY1 treatments was 0.368). Calcein intensity from corresponding to XY2 and XY3 treatments on neurons was in-between that for XY1 and XY4 treatments.

**FIGURE 3:**
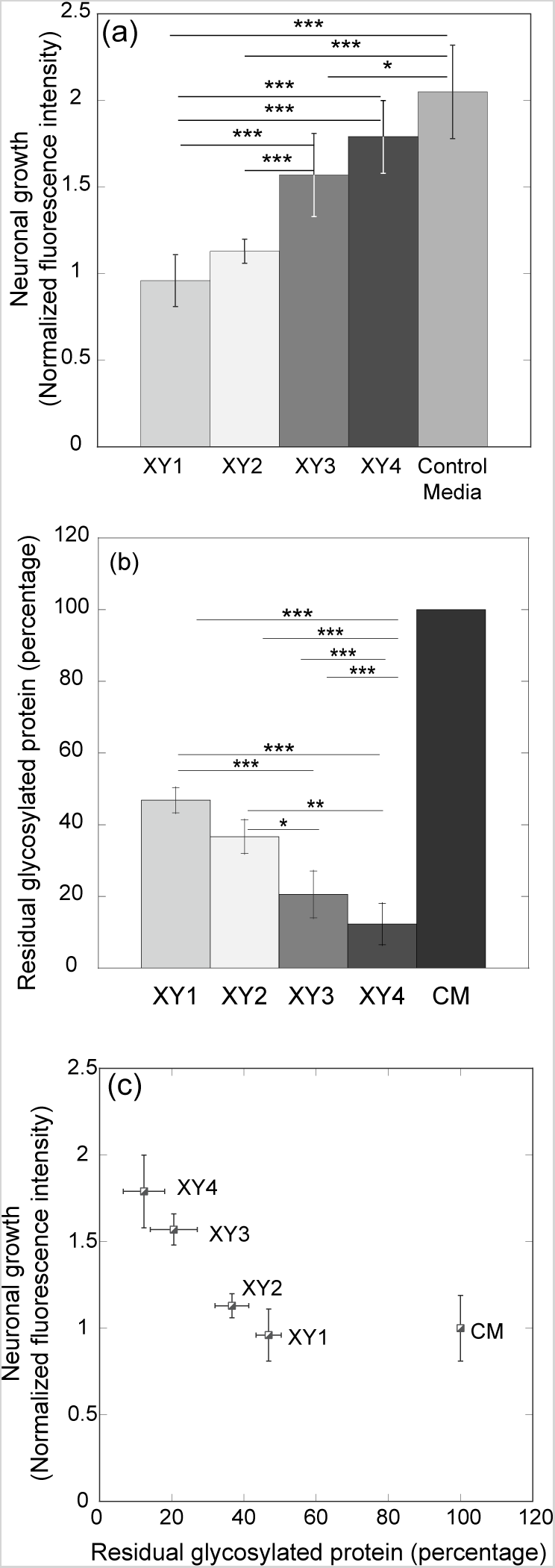
Comparison of neuronal outgrowth with extent of CSPG-under glycosylation. (a) Normalized fluorescence intensities from obtained from Calcein AM treatment on neurons. (b) Relative amount of CSPGs synthesized by astrocytes in response to different click-xylosides at 100 µM concentration. (c) Correlation of neuronal outgrowth and amount of CSPG present in CM. Each bar represents mean of three independent experiments. Error bar corresponds to standard deviation, *p < 0.01; **p < 0.001; ***p < 0.0001.

When the xylosides were directly added to neurons in fresh media, no difference in neurite growth compared to the untreated control was observed, suggesting that the click-xylosides in the conditioned media do not act directly on the neurons to promote growth (Fig S2). Instead, the click-xylosides likely acted on the astrocytes and altered the secreted contents of the CM. The molecular basis of varying neuronal outgrowth from click-xyloside treated CM could only be explained by characterizing the inhibitory CSPGs present in the CM. By labeling GAG chains secreted by click-xylosides treated astrocytes with radioactive [^35^S]Na_2_SO_4_, we determined the level of glycosylation, and sulfation patterns of the CS chains. As expected, click-xyloside treatment resulted in synthesis of protein-free, xyloside primed CS chains in the CM, as evident from the emergence of the second broad peak in SEC chromatograms (Fig S3). By multiplying the area percentages of the CSPG peaks with the total scintillation counts in purified CM, we calculated the relative amounts of glycosylated CSPGs present in the CM. As shown in Fig 3 (b), the reduction in the percentage of CSPGs present in the CM after click-xyloside treatment ranged from 63% (XY1) to 18% (XY4) of CSPGs secreted by untreated reactive astrocytes.

Sulfation patterns of CS chains influence their functionality. Therefore, we analyzed the sulfation profile of CS present in CM. As shown in Fig S4, CS chains derived from reactive astrocytes were mainly 4-*O*-sulfated, in agreement with earlier observation ^15^. Primed CS chains exhibited identical sulfation patterns with untreated reactive astrocytes; hence, the only significant difference among various click-xylosides-primed CM was the amount of glycosylated CSPGs and free CS chains present in the CM. Therefore, it is evident that the attachment of CS chains onto core proteins is critical to mediate neuronal inhibition. Fig 3 (c) compares neuronal growth with the relative amount of residual glycosylated CSPGs present in the media. CM containing higher amount of residual CSPGs results in lower neuronal growth and viability. Neuronal outgrowth is completely restored with 82% reduction in the CSPG content of reactive astrocytes. However, the inability of XY1-primed CM to rescue outgrowth suggests that even a close to 63% reduction in CSPGs did not reduce the inhibitory potential of astrocyte CM.

This study demonstrates that click-xylosides can modulate CSPG biosynthesis in astrocytes, which in turn contribute to neurite growth. Due to their enhanced bio-availability, click-xylosides prime GAG chains efficiently at lower concentrations in various cell lines^12^. Here, we studied the effect of click-xyloside treated astrocytes on neuronal growth. We showed that the outgrowth in CM-treated neurons correlates with the CSPG content present in the CM. From these results, we conclude that by modulating the CSPG biosynthesis in astrocytes, one can inhibit the neurotoxic effect and the inhibitory potential of the astrocyte-CM. In summary, click-xylosides, through inhibiting CS attachments to PGs and thereby resulting in the under-glycosylation of CSPGs, hold tremendous therapeutic potential in the treatment of CNS lesions and traumatic brain injury.

## Materials and Methods

### Materials

4-Methylumbelliferyl-β-D-xylopyranoside (4-MUX) was obtained from Sigma Aldrich (Cat # M7008). Click-xylosides were synthesized as reported earlier (Victor, X. V.; Nguyen, T. K. N.; Ethirajan, M.; Tran, V. M.; Nguyen, K. V.; Kuberan, B. *Journal of Biological Chemistry* 2009, *284*, 25842). Embryonic day-18 primary rat hippocampal cells (Cat# PC35101) were purchased from Neuromics Inc. Calcein AM was from Cultrex (Cat#: 4892-010-01).

### Primary rat astrocyte preparation

Primary rat cortical astrocytes were isolated from P1 Sprague-Dawley rats in accordance with the protocol approved by University of Utah Institutional Animal Care and Use Committee (McCarthy, K. D.; De Vellis, J. *The Journal of cell biology* 1980, *85*, 890). Briefly, meninges were removed and cortical tissues were broken and digested in 1.33% collagenase for 30 min. Cortices were then treated with 0.25% trypsin for 30 min, triturated, suspended, and plated in T75 tissue culture flasks. Astrocytes were purified by shaking the culture overnight at 175 rpm, for one week postdissection. After shaking, cells were dissociated with 0.25% trypsin-EDTA and either frozen in liquid N_2_ or prepared for use. Cells were seeded onto six-well tissue culture plates at a density of 10,000 cells cm^-2^. Astrocytes were maintained in DMEM/F12 (Gibco) supplemented with 10% fetal bovine serum (FBS, Sigma) until they were confluent. Upon reaching confluence, the media was replaced with F12 supplemented with 10% dialyzed FBS (Sigma) for xyloside treatment and S-35 labeling.

### Neuronal culture

Embryonic day-18 primary rat hippocampal cells (Cat# PC 35101) were purchased from Neuromics Inc. Dissociated hippocampal neurons were pelleted and suspended in NbActiv1 media (Catalog # NBActiv1-100, BrainBits) to a concentration of 8 x 10^4^ cells/ml. Two milliliters of cell suspension was placed into each well of a six-well plate (containing poly-L-lysine coated coverslips). In some cases, 100 µl of cell suspension was plated into each well of a pLL-coated 96-well plate. Neurons were maintained at 37ºC and 5% CO_2_. After reaching the desired time point, cells were fixed and stained with DAPI and chicken anti-rat TAU/MAPT primary antibody (Neuromics, CH22113), and labeled with goat anti-rat IgG secondary antibody conjugated to AlexaFluor 488 (Molecular Probes, A11039).

### Imaging

Fluorescent images were collected with an Olympus FV1000 IX81 microscope using a Planapo oil immersion 20x objective, NA 0.75. The confocal aperture was set to one airy unit and these settings were maintained between comparable samples. DAPI and AlexaFlour 488 were excited using 405 nm and 488 nm light, respectively. Acquisition software used was FVASW V 2.1.

### Calcein-based viability assay

Cell viability and neurite outgrowth was quantified using a Calcein AM stain. Briefly, the media from each well of a 96-well plate was removed and 100 µl of 2 µM Calcein AM solution was added. The plate was incubated for 30 min at 37ºC and 5% CO_2_. The viability was measured using a well plate reader by recording the absorbance at 485 nm excitation and 520 nm emission. Cells were also imaged using a fluorescence microscope. Each plate consisted of at least five replicates of corresponding treatments. Statistical analysis on fluorescence intensities was done by Anova (Kaleidagraph, Synergy Software).

### Radiolabelling of astrocyte-derived PGs

Astrocyte-derived PGs were radiolabelled to analyze the GAGs present in conditioned media (CM). The protocol for radiolabelling and purifying GAGs is described elsewhere ^2^. Briefly, astrocytes were plated in six-well plates and were maintained in F12 medium with 10% dialyzed fetal bovine serum. [^35^S]Na_2_SO_4_ (PerkinElmer) was added to media for radiolabeling of CS chains (10 µCi per ml).

### GAG purification

Conditioned media (CM) was collected from wells and was diluted with equal amount of water. The diluted CM was loaded on a 0.2 ml DEAE-Sepharose column pre-equilibrated with 10 column volumes of wash buffer (0.1M NaCl, 20mM NaOAc, 0.01% TritonX, pH 6.0). The column was then washed with 15 column volumes of wash buffer. Bound GAGs were eluted with 9 column volumes of elution buffer (1M NaCl, 20mM NaOAc, 0.01% TritonX, pH 6.8). The amount of GAGs contained in the CM was quantified by the ^35^S-radioactivity incorporated in the purified GAG. One microliter of eluate was diluted in 5 ml of scintillation mixture (PerkinElmer) and the amount of GAG was quantified using a scintillation counter for total radioactivity.

### Analysis of CS chains

The size of radiolabelled CS chains in the form of CSPGs or xylosides-primed CS chains was determined by measuring their migration time on a size exclusion column using high-pressure liquid chromatography (SEC-HPLC) with an inline flow scintillation analyzer. Samples containing 1 million total count were loaded on two G3000SW_XL_ columns (Tosoh, 7.8 mm x 30 cm), which were connected serially and eluted over 80 min with phosphate buffer (50mM mM NaH(PO_4_)_2_, 50mM NaH_2_(PO_4_), and 150mM NaCl, pH 6). The sulfation pattern of CS chains was determined by measuring the migration time of CS disaccharides on an anion-exchange column using HPLC with inline flow scintillation analyzer. CS disaccharides were obtained by digesting CM with Chondroitinase ABC (ChABC, 5 mU, Seikagaku Corp.) in 10 µl of buffer (70 mM CH_3_COONa; 0.2% BSA, pH 6.0). The mixture was incubated for 12 hr at 37 ºC. Resulting disaccharides were then analyzed using a strong anion-exchange (SAX)-HPLC column coupled to a UV detector. The disaccharides were eluted at a flow rate of 1 ml/min and resolved with a linear gradient of 0% to 100% solvent B (800 mM NaH_2_PO_4_) in solvent A (100 mM NaH_2_PO_4_) over 40 min. CS disaccharide standards (Seikagaku Corp. Cat # 400571) were used for characterizing the disaccharide composition of CS variants.

## Ackowledgements

The study was supported by NHLBI sponsored Programs of Excellence in Glycosciences grant, HL107152. We thank M. Kalita, M.V. Quintero, V.M. Tran and R. Lin for synthesizing xylsoides, J. S. Chua for experimental verification and analysis, and V. Hlady and P. Tresco for providing astrocytes and discussion. No competing financial interests have been declared. V. S. published the described work herein as part of his doctoral thesis work.

---

**Figure S1.**
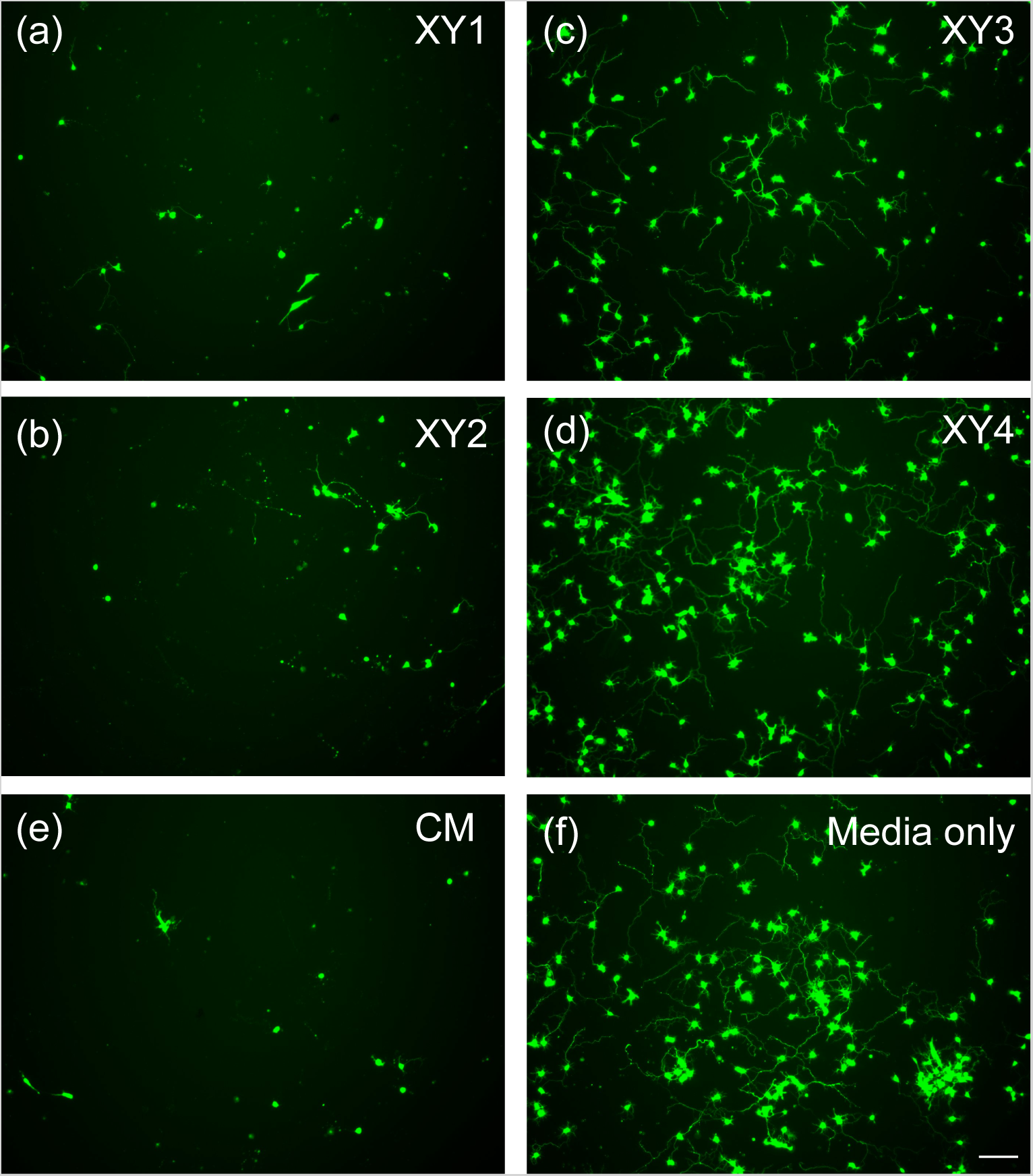
Neuronal response to mono-xylosides, evident from Calcein AM staining. (a), (b), (c) and (d) represent images of neurons after XY1, XY2, XY3 and XY4 based growth assay. (e) and (f) are neurons treated with control CM and fresh media respectively. (Scale bar = 100 μm, green: Calcein AM)

**Figure S2.**
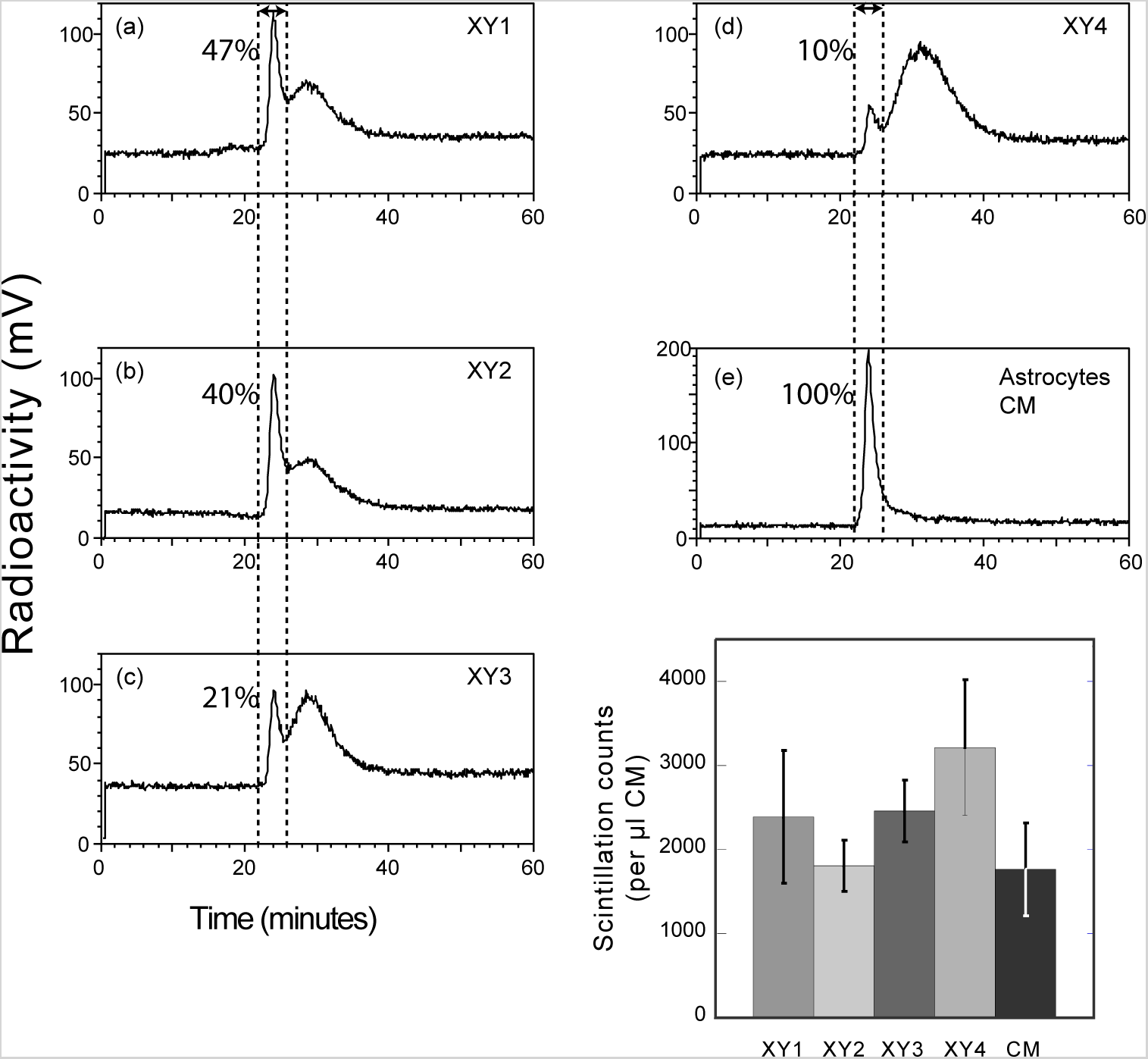
The elution profiles of CS chains primed by mono-xyloside through a SEC column. (a-d) Elution profiles of CS present in astrocyte CM after treatment with XY1, XY2, XY3, and XY4, respectively. (e) The profile of CS secreted by control astrocytes. As compared to control astrocytes, click-xyloside treatments show a second peak at later time point, corresponding to free CS chains primed by exogenously added click-xylosides. The first peak (21 – 23 min, indicated by the dotted lines) corresponds to CSPGs synthesized endogenously by the cells. The area percentage of each CSPG peak is specified in each figure. (f) The scintillation counts of 1 μl aliquot of purified CM (indicating the total amount of CS chains present in the media). The CSPG area percentage was multiplied by the scintillation counts from purified CM to obtain the amount of CSPG present in the CM. Each bar represents mean of three independent experiments. Error bar denotes standard deviation.

**Figure S3.**
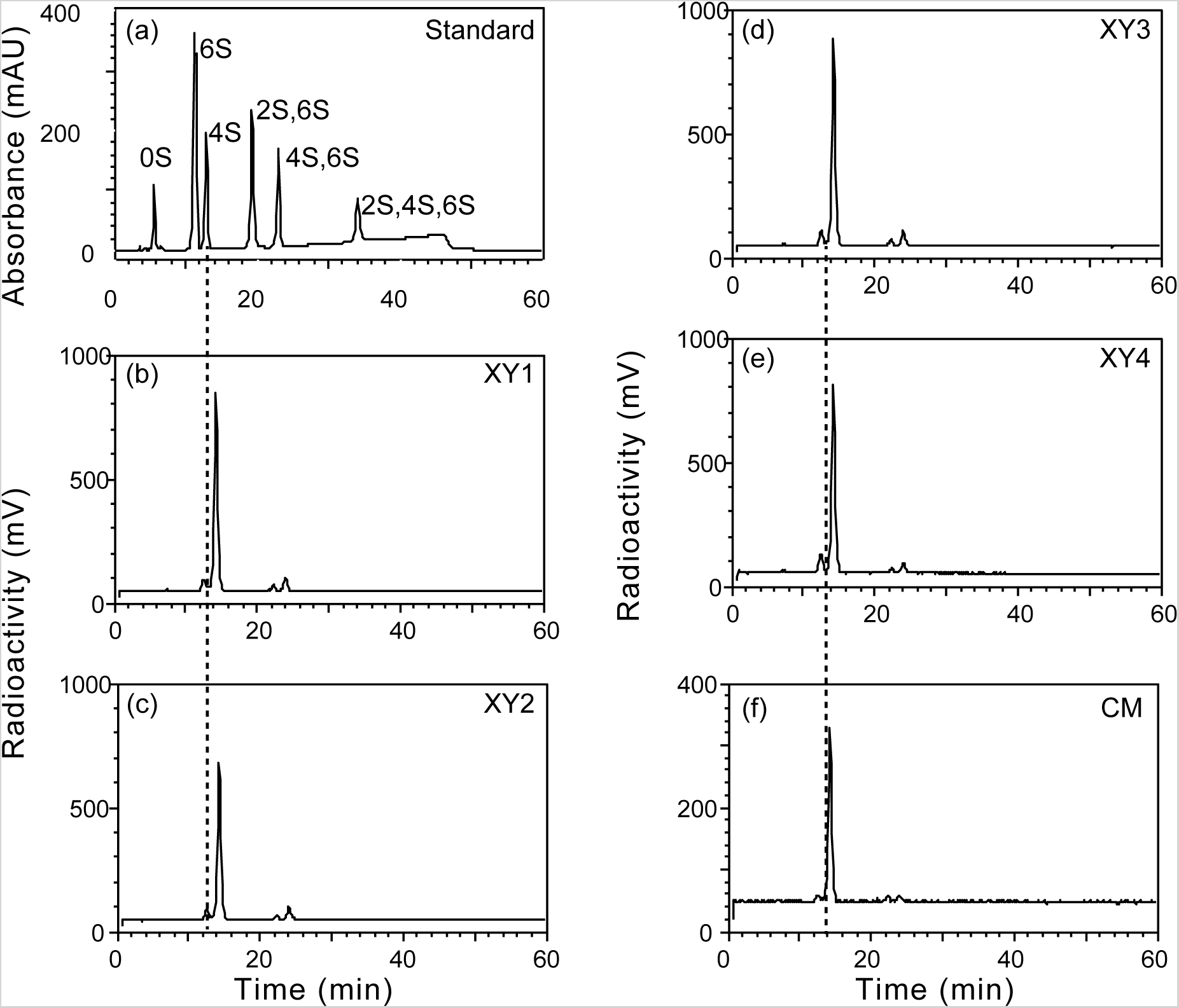
Sulfation profiles of CS chains obtained by passing the disaccharide samples through a strong anion exchange HPLC. (a) Elution profile of CS disaccharide standards created by measuring the UV absorbance of the disaccharides. (b-e) Sulfation profiles of CS present in astrocyte CM after treatment with XY1, XY2, XY3, and XY4, respectively. (e) The profile of CS secreted by control astrocytes. The flow time of one minute between the UV detector and the radiometer caused the radiolabelled disaccharide peak to lag behind the corresponding disaccharide standards (indicated by the dashed line).

